# Distinct steady-state properties and TNF responses of epithelial long myosin light chain kinase (MLCK) splice variants

**DOI:** 10.1101/2022.10.13.512159

**Authors:** Sandra D. Chanez-Paredes, Shabnam Abtahi, Juanmin Zha, Li Zuo, Weiqi He, Jerrold R. Turner

**Affiliations:** Laboratory of Mucosal Barrier Pathobiology, Department of Pathology, Brigham and Women’s Hospital and Harvard Medical School, Boston, MA, USA; Jiangsu Key Laboratory of Neuropsychiatric Diseases and Cambridge-Suda (CAM-SU) Genomic Resource Center, Soochow University, and Department of Oncology, The First Affiliated Hospital of Soochow University, Suzhou, China; Anhui Medical University, Hefei, Anhui, China, 230032

**Author notes:** Correspondence: Weiqi He and Jerrold R. Turner.

**Keywords:** actin, intestinal epithelium, tight junction, inflammatory bowel disease (IBD), tumor necrosis factor (TNF), myosin light chain kinase, intestinal permeability, barrier dysfunction

## Abstract

Intestinal epithelia express two long myosin light chain kinase (MLCK) splice variants, MLCK1 and MLCK2. Unlike MLCK2, MLCK1 is concentrated at the perijunctional actomyosin ring and this localization is enhanced by tumor necrosis factor (TNF) signaling. Here we sought to identify and characterize the domain(s) that direct basal and TNF-induced MLCK1 subcellular localization. Quantitative morphometry demonstrated specific increases in MLCK1 expression and perijunctional localization in Crohn’s disease patient biopsies, relative to controls. TNF induced perijunctional recruitment of MLCK1-EGFP but did not affect localization of MLCK2-EGFP, which was predominantly associated with basal stress fibers. Recombinant N-terminal MLCK1 and MLCK2 regions accelerated actin polymerization in vitro but were not different from one another. In contrast, the affinity of N-terminal MLCK1 binding to F-actin was greater than that of MLCK2. Perijunctional MLCK1 and MLCK2 domain recruitment in intestinal epithelial cells paralleled in vitro F-actin binding. The unique MLCK1 Ig3 domain was necessary, but not sufficient, for both F-actin binding and perijunctional recruitment, but, nevertheless, displaced perijunctional MLCK1, enhanced steady-state barrier function, and limited TNF-induced MLCK1 recruitment and barrier loss. These data demonstrate selective perijunctional MLCK1 recruitment in Crohn’s disease, suggest that F-actin binding contributes to perijunctional recruitment, and show that Ig3 can act as a dominant negative effector that limits TNF-induced MLCK1 recruitment and barrier loss. These results data provide key mechanistic detail that will enable development of therapeutics that target Ig3, or its intercellular binding partners, to reverse inflammation-induced barrier loss and limit disease progression.

## INTRODUCTION

The critical role of MLCK in physiologic and pathologic regulation of intestinal epithelial tight junctions was first reported over 20 years ago (1,2) and, since that time, has been confirmed using in vitro and in vivo models, and in diseased human tissues (3–16). The initial cloning of human intestinal epithelial MLCK identified two splice variants of long MLCK and showed that short, smooth muscle, MLCK is not expressed within human intestinal epithelia. MLCK1 and MLCK2 splice variants differ only by the removal of exon 11, composed of 207 nucleotides, from the latter. As a result, the Ig3 domain within MLCK2 is incomplete. MLCK1 and MLCK2 also display distinct, differentiation-dependent expression patterns, with MLCK1 was limited to villus epithelia and MLCK2 expressed throughout the crypt-villus axis (17).

We recently demonstrated that inflammatory stimuli, including tumor necrosis factor (TNF), trigger recruitment of MLCK1, but not MLCK2, recruitment to the perijunctional actomyosin ring in vitro and in vivo (6,18). Because primary sequence within Ig3 is the only structural difference between these long MLCK splice variants, we reasoned that this region must be essential for perijunctional MLCK1 recruitment. Our previous work also shows that targeting MLCK1 Ig3 domain or its binding partner, FKBP8, can displace MLCK1 from the perijunctional actomyosin ring, prevent or reverse TNF-induced barrier loss in vitro and in vivo, and limit progression of experimental inflammatory bowel disease (IBD) in mice (6,18). Nevertheless, the contributions of Ig3 to MLCK1 regulation and function are poorly understood and it is not clear if, for example, this domain alone is sufficient to direct perijunctional MLCK1 recruitment.

Here, we show that Ig3 alone is incompetent for F-actin binding in vitro and insufficient to direct MLCK1 recruitment within intestinal epithelial cells. Although Ig3 is distributed diffusely throughout the cytoplasm, it can function as a dominant negative effector that prevents steady-state and TNF-induced MLCK1 recruitment and limits TNF-induced barrier loss. These data provide mechanistic insight into the mechanisms that direct MLCK1 subcellular localization as well as means of disrupting these processes. The resulting insight will provide a foundation for development of molecular therapies that interfere with MLCK1 recruitment and prevent inflammation-associated intestinal barrier loss and disease progression.

## RESULTS

### MLCK1 is upregulated and specifically recruited to the perijunctional actomyosin ring in Crohn’s disease

We recently reported that, similar to well-differentiated Caco-2_BBe_ monolayers, TNF increased long MLCK expression and perijunctional MLCK1 recruitment in human duodenal enteroids (8,18,19). To determine if this is representative of human disease, we analyzed ileal biopsies from Crohn’s disease patients and healthy controls by quantitative immunofluorescence using MLCK1-specific or pan-long MLCK antibodies. MLCK1 was primarily expressed in villus epithelium and concentrated at the perijunctional actomyosin ring and within the apical cytoplasm in biopsies from control subjects (Fig. 1A, B). Total long MLCK was present throughout the cytoplasm and on epithelial lateral membranes throughout the crypt-villus axis but only a minor component was present at the perijunctional actomyosin ring (Fig. 1A, B).

**Figure 1.**
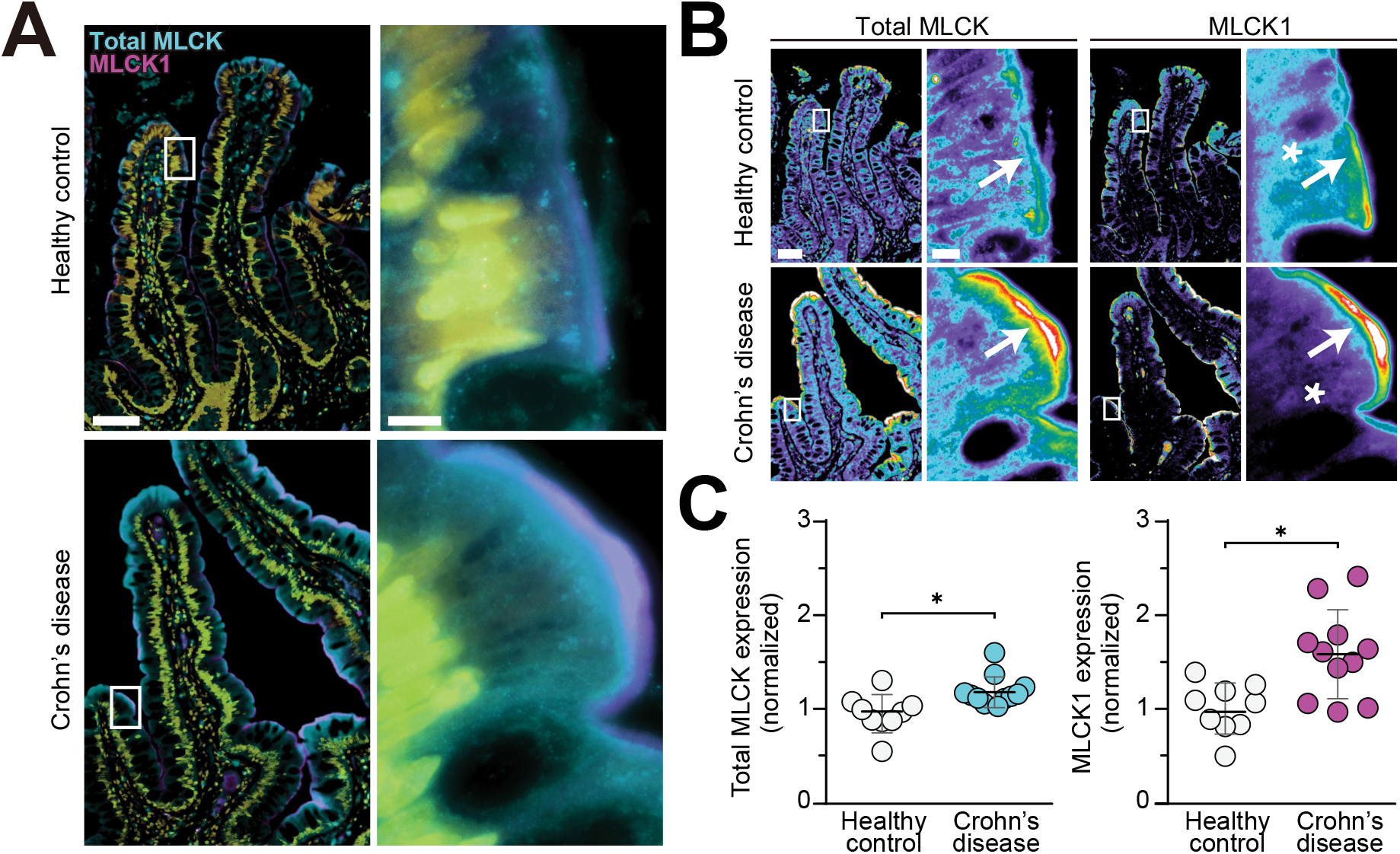
MLCK1 expression and recruitment to the perijunctional actomyosin are specifically increased in Crohn’s disease. **A.** Representative immunostains of ileal biopsies from Crohn’s disease patients and healthy controls labeled for long MLCK1 (magenta), total long MLCK (cyan), and nuclei (yellow). Note the dense MLCK1 band at the perijunctional actomyosin ring in Crohn’s disease. In contrast, the perijunctional actomyosin ring contains relatively little MLCK1 in healthy control tissues. Bars = 50 μm, 5 μm. **B.** Total MLCK or MLCK1 pseudocolor heatmap images of the same fields as A. The small amount of MLCK1 expressed in healthy controls is present at the perijunctional actomyosin ring (arrow) and within the cytoplasm (asterisk). In contrast, MLCK1 expression in Crohn’s disease is almost exclusively within the perijunctional actomyosin ring (arrow), and there is little cytoplasmic staining (asterisk). Note the overall increase in total MLCK expression and perijunctional localization in Crohn’s disease. Bars = 50 μm, 5 μm. **C.** Quantitative morphometry of total MLCK and MLCK1 staining in individual biopsies. Each point represents a distinct patient. *n* = 9 healthy controls, *n* = 11 Crohn’s disease patients. Mean ± SD. Student’s *t* test. *, P < 0.05.

MLCK1 and total MLCK expression were both significantly increased throughout the crypt-villus axis in Crohn’s disease (Fig. 1B, C), where MLCK1 was almost entirely restricted to the perijunctional actomyosin ring (Fig. 1B). Moreover, the magnitude of increased MLCK1 expression in Crohn’s disease was significantly greater than that of total MLCK (Fig. 1C), suggesting that inflammatory stimuli bias mRNA splicing towards MLCK1 translation. Total long MLCK expression, MLCK1 expression, and MLCK1 recruitment to the perijunctional actomyosin ring are, therefore, all increased in Crohn’s disease.

### MLCK1 and MLCK2 differentially associate with distinct actin-rich structures

Although we can infer that structures labeled with anti-total MLCK but not anti-MLCK1 reflect MLCK2 localization, the absence of unique sequences within MLCK2 makes it impossible to develop MLCK2-specific antisera. To overcome this, we stably expressed MLCK1-EGFP and MLCK2-EGFP and assessed their subcellular localization in Caco-2_BBe_ cell monolayers. Western blot using anti-EGFP or anti-pan-long MLCK antibodies demonstrated equivalent expression of MLCK1-EGFP or MLCK2-EGFP in independent transfected populations (Fig. 2A). Nevertheless, MLCK1-EGFP and MLCK2-EGFP display distinct intracellular distributions. MLCK1 was concentrated within the perijunctional actomyosin ring, where it colocalized with ZO-1 and, to a lesser degree, along lateral membranes (Fig. 2B, D, E). In contrast, MLCK2 was prominently associated with basal stress fibers and was also present at lateral membranes (Fig. 2B-F). Both the localization and functional impact of MLCK2 expression are, therefore, distinct from those of MLCK1.

**Figure 2.**
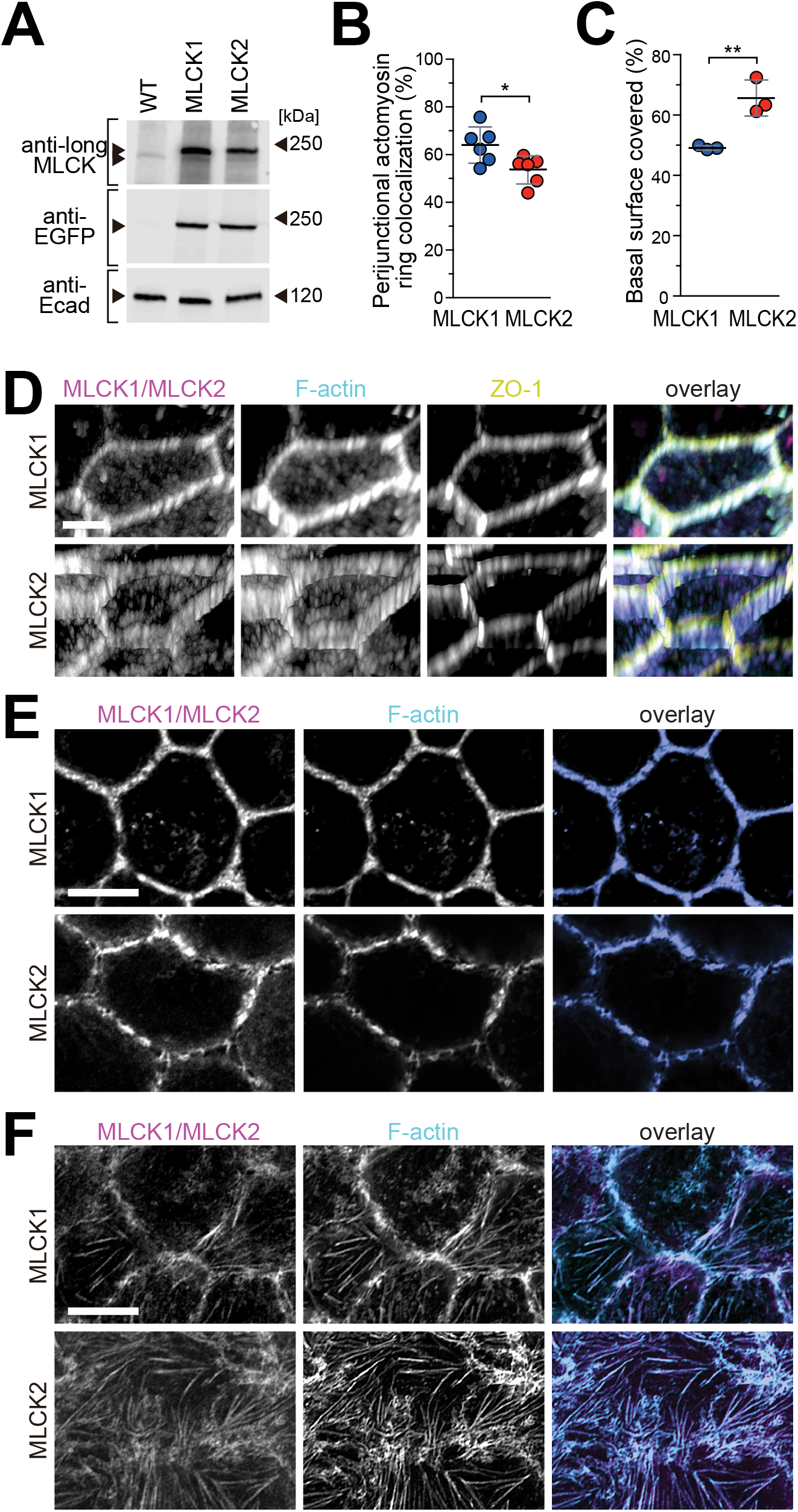
MLCK1 and MLCK2 have distinct intracellular distributions. **A.** Western blot analysis of lysates of wild type (WT) or stably-transfected Caco-2_BBe_ monolayers. Long MLCK antibody labeling detects endogenous MLCK (lower arrowhead) as well as MLCK1-EGFP and MLCK2-EGFP (upper arrowhead). Anti-EGFP detects only MLCK1-EGFP and MLCK2-EGFP. E-cadherin (Ecad) is shown as a loading control. MLCK1-EGFP and MLCK2-EGFP are expressed at similar levels to one another. Both are overexpressed relative to endogenous long MLCK. **B.** MLCK1-EGFP (blue circles) is preferentially recruited to the perijunctional actomyosin ring, relative to MLCK2-EGFP (red circles). *n* = 6. Mean ± SD. Student’s *t* test. *, P < 0.05. **C.** MLCK2-EGFP is concentrated at the basal surface, where it colocalizes with F-actin stress fibers. *n* = 3. Mean ± SD. Student’s *t* test. **, P < 0.01. **D.** 3D projections showing subcellular localization of MLCK1-EGFP (magenta, upper row) or MLCK2-EGFP (magenta, lower row) as well as F-actin (cyan) and ZO-1 (yellow). Note the colocalization of MLCK1-EGFP with ZO-1. In contrast, MLCK2-EGFP is present at lateral membranes, below the tight junction, and at the basal surface. Bar = 2.5 μm. **E.** Confocal slices at the level of the perijunctional actomyosin ring demonstrate a dense, continuous, ring of MLCK1-EGFP with little cytoplasmic signal. In contrast, perijunctional MLCK2-EGFP is less dense, discontinuous, and includes faint staining adjacent to the perijunctional actomyosin ring. Bar = 5 μm. **F.** Confocal slices at the level of the basal stress fibers show colocalization MLCK2-EGFP with F-actin stress fibers. MLCK1-EGFP also associated with stress fibers but is more prominent at lateral membranes. Bar = 5 μm.

### Subcellular localizations of MLCK1 and MLCK2 are determined separately and are independent of kinase activity

The subcellular distributions of MLCK1 and MLCK2 were each consistent within several monoclonal lines as well as polyclonal populations of transfected Caco-2_BBe_ cells. Nevertheless, we considered the possibility that the unique splice variant distributions could be due to clonal variation rather than true biological differences. Moreover, we asked whether the distinct subcellular localizations of MLCK1 and MLCK2 modified by one another. Finally, although the MLCK catalytic domain is at the C-terminal end of the molecule, far from the N-terminal sequences that, in MLCK1, complete Ig3, we asked whether enzymatic activity was necessary for correct MLCK1 and MLCK2 localization. In order to address the first two questions, we co-expressed MLCK1-fusion red (FR) and MLCK2-EGFP in Caco-2_BBe_ cells. Both splice variants were associated with F-actin-rich structures and, similar to cells expressing only one fluorescent-tagged splice variant, MLCK1 was primarily associated with the perijunctional actomyosin ring and microvilli while MLCK2 was more strongly associated with basal stress fibers and lateral membranes (Fig. 3A, B, D). Thus, clonal variation does not explain the differential subcellular localizations of MLCK1-EGFP and MLCK2-EGFP. More importantly, the distributions of these splice variants are not affected by one another. Finally, despite limited qualitative effects on F-actin, including reduced prominence of basal stress fibers, MLCK enzymatic inhibition with the highly-specific and stable pseudo-substrate peptide Dreverse PIK (1,20,21) did not affect MLCK1-FR and MLCK2-EGFP subcellular distributions (Fig. 3B, C, D).

**Figure 3.**
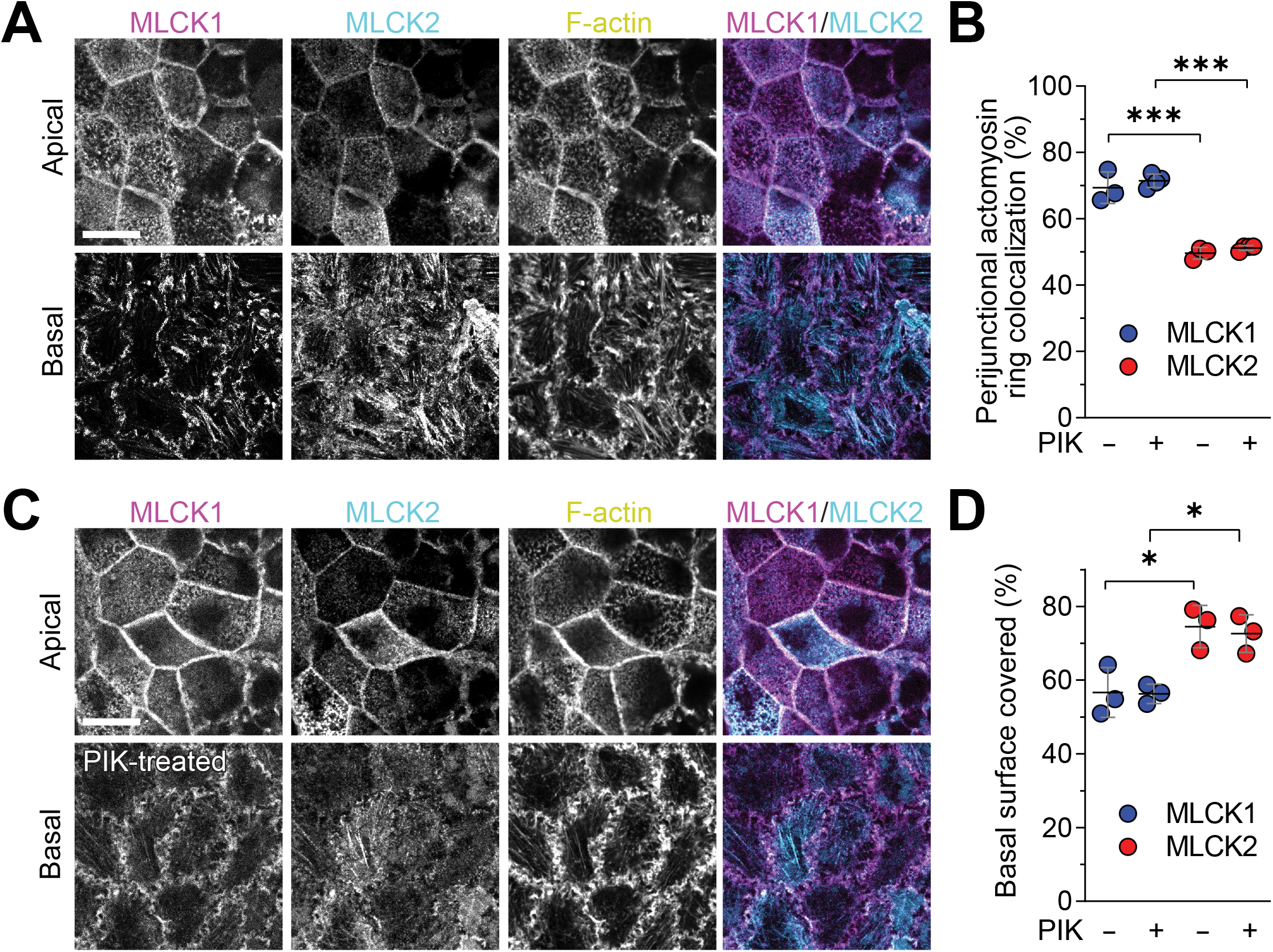
MLCK1 and MLCK2 distributions are independent of their kinase activity. **A.** Confocal slices at the level of the perijunctional actomyosin ring (apical) or basal stress fibers (basal) in cells co-expressing both MLCK1-FR and MLCK2-EGFP demonstrate greater MLCK1 localization of the perijunctional actomyosin ring, relative to MLCK2. In contrast, MLCK2 decorates basal stress fibers more extensively than MLCK1, which is also present at lateral membranes. All images are from a single z-stack. Bar = 10 μm. **B.** Quantitative morphometry confirms greater MLCK1 recruitment to the perijunctional actomyosin ring, relative to MLCK2. *n* = 3-4. Mean ± SD. ANOVA. ***, P <0.001. **C.** Inhibition of MLCK enzymatic activity using the highly-specific MLCK inhibitor, PIK (250 μM, 2 hours), did not distributions of MLCK1 or MLCK2. All images are from a single z-stack. Bar = 10 μm. **D.** Quantitative morphometry confirms preferential association of MLCK2 with basal stress fibers, relative to MLCK1. *n* = 3. Mean ± SD. ANOVA. *, P < 0.05.

### MLCK1 interactions with the cytoskeleton are more stable than MLCK2 interactions

We and others have shown that MLCK-mediated MLC phosphorylation promotes F-actin polymerization (22–25) and that the enhanced F-actin polymerization induced by this regulated MLC activation can limit epithelial migration (26,27). Given the striking differences in basal stress fiber density induced by MLCK2 overexpression, we asked if this correlated with changes in epithelial migration. Migration of MLCK1-EGFP-expressing Caco-2_BBe_ cells was indistinguishable from that of non-transfected cells. Conversely, MLCK2-EGFP expression markedly inhibited Caco-2_BBe_ cell migration (Fig. 4A). Together with preferential localization along stress fibers, this result suggests that, in contrast to MLCK1, the primary function of MLCK2 may be as a cytoskeletal effector during epithelial migration.

**Figure 4.**
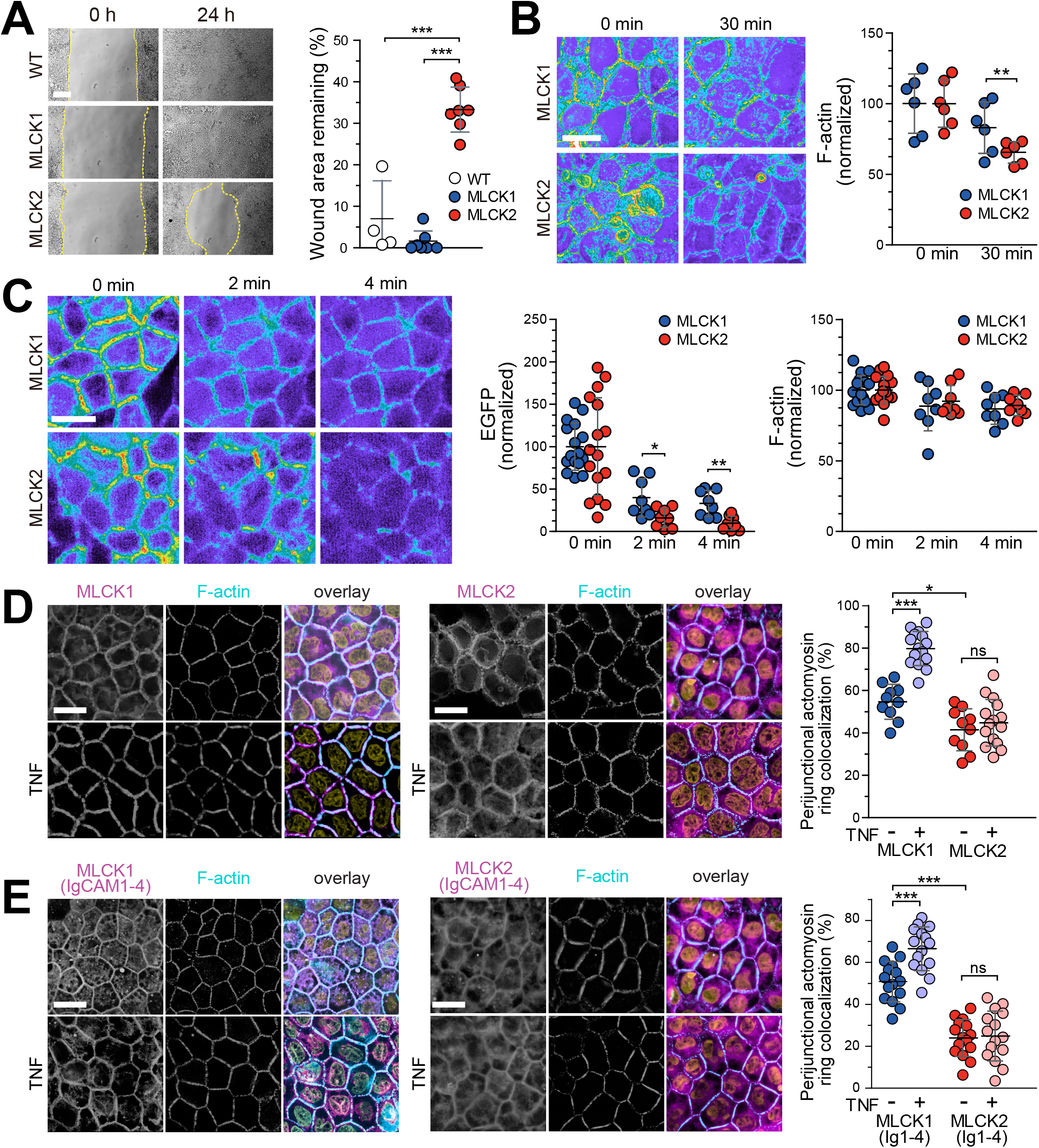
Divergent MLCK1 and MLCK2 behaviors. **A.** MLCK1-EGFP expression does not impact epithelial migration relative to wild type (WT), non-transfected cells. In contrast, MLCK2-EGFP expression markedly delays migration and wound closure. Representative images are shown. Bar = 100 μm. *n* = 4-7. Mean ± SD. ANOVA. ***, P < 0.001. **B.** Representative pseudocolor heatmap images show F-actin labeled before or 30 min after addition of LatA (1 μM) to monolayers expressing MLCK1-EGFP or MLCK2-EGFP. F-actin content was similar before LatA treatment. F-actin was resistant to LatA in MLCK-EGFP-expressing cells, relative to MLCK2-EGFP-expressing cells. Bar = 10 μm. *n* = 6. Mean ± SD. Student’s *t* test. **, P < 0.01. **C.** Monolayers expressing MLCK1-EGFP or MLCK2-EGFP were treated with detergent-containing extraction buffer. Representative pseudocolor heatmap images collected at 2 minute intervals are shown. MLCK1-EGFP retention was greater than that of MLCK2-EGFP. Representative samples were fixed at each time point and labelled for F-actin, which was similar in MLCK1-EGFP- and MLCK2-EGFP-expressing monolayers. Bar = 10 μm. *n* = 8 - 16. Mean ± SD. ANOVA. *, P < 0.05. **, P < 0.01. **D.** Caco-2_BBe_ monolayers with stable expression of EGFP-tagged MLCK1, MLCK2, MLCK1(Ig1-4), or MLCK2(Ig1-4) were treated with TNF or vehicle, as indicated, for 4 hours. EGFP-tagged constructs (magenta), F-actin (cyan), and nuclei (yellow) are shown in the overlay images. Morphometric analyses shows that recruitment of MLCK1 and MLCK2 Ig1-4 truncation mutants is similar to their corresponding full-length MLCK constructs. Bars = 10 μm. *n* = 10 - 15. Mean ± SD. ANOVA. *, P <0.05. ***, P <0.001.

Previous studies have shown that the affinity of MLCK1 for F-actin and myofilaments is greater than that of short, smooth muscle MLCK and that this correlated with MLCK1-mediated stabilization of F-actin structures (28). We therefore asked whether MLCK1 and MLCK2 differed in their abilities to stabilize F-actin. To test this, monolayers were treated with latrunculin A (LatA), which, as expected, reduced F-actin content (Fig. 4B). MLCK1-EGFP-expressing monolayers were, however, relatively protected and retained significantly more F-actin than MLCK2-EGFP-expressing monolayers (Fig. 4B). These data suggest that MLCK1 may stabilize actin-based structures more effectively than MLCK2 and may, therefore, also explain the relative resistance of perijunctional actin to drug-induced depolymerization (29).

To determine if mediated microfilament stabilization was associated with increased MLCK1 binding, monolayers were imaged following addition of a detergent-containing cytosolic extraction buffer. Although MLCK1-EGFP and MLCK2-EGFP were both released after extraction, MLCK2-EGFP loss was significantly greater than that of MLCK1-EGFP (Fig. 4C). We considered the possibility that this could be related to microfilament stabilization by MLCK1. However, the observation that F-actin content was not significantly reduced by extraction (Fig. 4C) excludes this mechanism. Taken together with the distinct effects of MLCK1 and MLCK2 on LatA-induced F-actin depolymerization, these data suggest that interactions between microfilaments and MLCK1 are more stable than those with MLCK2.

### The N-terminal extension that distinguishes long MLCK isoforms from smooth muscle MLCK is sufficient to direct MLCK1, but not MLCK2, to the perijunctional actomyosin ring

The short 69 amino acid sequence encoded by exon 11 is the only difference between MLCK1 and MLCK2. This sequence completes Ig3 and has been suggested to be a substrate for Src kinase-mediated phosphorylation-induced increases in enzymatic capacity (6,30). We therefore considered the hypothesis that differences subcellular localizations of MLCK1 and MLCK2 are, in part, mediated by the kinase domain. To test this, we compared distributions of full-length MLCK1 and MLCK2 to those of corresponding truncation mutants that lacked the catalytic domain and consisted only of Ig1-4 domains. Similar to full-length MLCK1-EGFP, MLCK1(Ig1-4)-EGFP was concentrated at the perijunctional actomyosin ring at steady-state and recruited to the perijunctional actomyosin ring following TNF treatment (Fig. 4D, E). In contrast, neither MLCK2-EGFP nor MLCK2(Ig1-4)-EGFP were concentrated at the perijunctional actomyosin ring or redistributed in response to TNF (Fig. 4D, E). These data exclude an essential role of the MLCK catalytic domain as a regulator of subcellular distribution and demonstrate that Ig1-4 MLCK1 is sufficient to direct both basal and TNF-induced recruitment to the perijunctional actomyosin ring.

### MLCK1(Ig1-4) affinity for F-actin is greater than that of MLCK2(Ig1-4)

Previous work has identified an actin-binding region within the first two Ig domains of long (31). These are identical in MLCK1 and MLCK2 and cannot, therefore, explain the preferential recruitment of MLCK1 and MLCK1(Ig1-4) to the perijunctional actomyosin ring. We therefore assessed MLCK1(Ig1-4) and MLCK2(Ig1-4) binding to actin during microfilament polymerization. MLCK1(Ig1-4) and MLCK2(Ig1-4) were equally soluble in the absence of actin (Fig. 5A). In the presence of G-actin, MLCK1(Ig1-4) and MLCK2(Ig1-4) each enhanced F-actin polymerization, relative to BSA, and co-sedimented with the F-actin pellet similarly (Fig. 5A, B). Although both MLCK1(Ig1-4) and MLCK2(Ig1-4) enhanced the extent of G-actin polymerization at completion, we considered the possibility that they might do so with distinct kinetics. However, kinetic analyses showed that MLCK1(Ig1-4) and MLCK2(Ig1-4) accelerated the rate and increased the extent of actin polymerization similarly (Fig. 5C). The distinct subcellular localizations, effects on LatA-induced actin depolymerization, and cytoskeletal retention during detergent extraction of MLCK1 and MLCK2 cannot, therefore, be explained by differences between MLCK1(Ig1-4) and MLCK2(Ig1-4) in vitro effects on actin polymerization.

**Figure 5.**
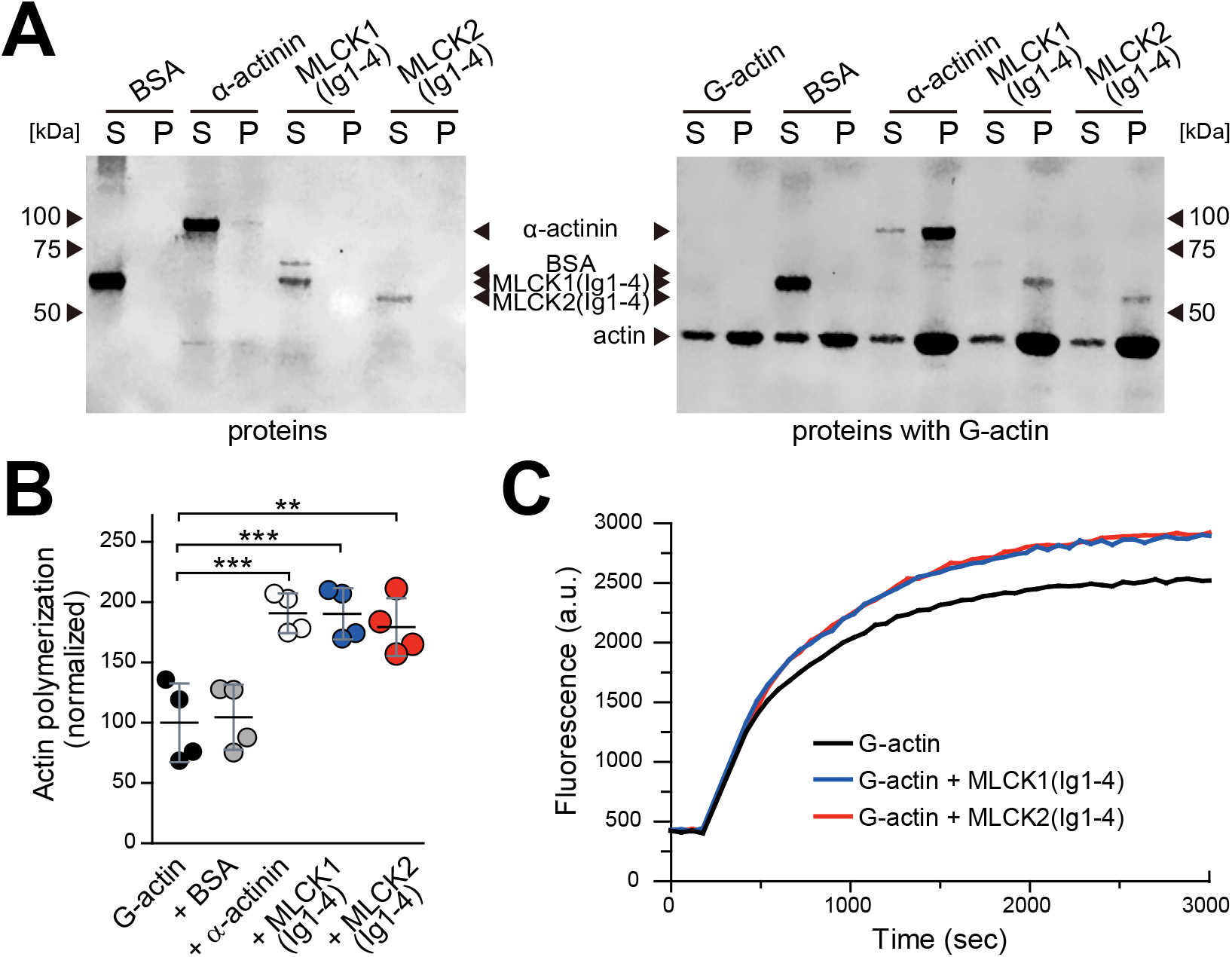
MLCK1(Ig1-4) and MLCK2(Ig1-4) promote actin polymerization. **A.** Spin-down assays were performed to assess F-actin binding and polymerization enhancement activity. In the absence actin (left), bovine serum albumin (BSA, 66 kDa), α-actinin (100 kDa), MLCK1(Ig1-4) (67 kDa), and MLCK2(Ig1-4) (59 kDa) all failed to sediment in the pellet fraction (P) and were entirely within the supernatant (S) fraction. When pre-incubated with G-actin (right), α-actinin, MLCK1(Ig1-4), and MLCK2(Ig1-4) all co-sedimented with the F-actin pellet. In addition, each enhanced actin polymerization relative to G-actin alone or BSA. In contrast to actin-binding proteins, BSA remained in the supernatant. **B.** Quantitative analysis showing extent of actin polymerization in the presence of α-actinin, MLCK1(Ig1-4), MLCK2(Ig1-4), or BSA. *n* = 4 Mean ± SD. ANOVA. **, P <0.01. ***, P <0.001. **C.** Pyrene-labeled G-actin polymerization was monitored by fluorescence. Both MLCK1(Ig1-4) and MLCK2(Ig1-4) accelerated the extent of polymerization. Traces shown are representative of *n* = 3 in this experiment.

We also considered the possibility that the MLCK1(Ig1-4) and MLCK2(Ig1-4) might differ in their affinities for F-actin. To test this, we analyzed the binding of recombinant MLCK1(Ig1-4), MLCK2(Ig1-4), MLCK1(Ig1-3), and MLCK1(Ig3) to F-actin using microscale thermophoresis (Fig. 6A). The Kd of MLCK1(Ig1-4) for F-actin was nearly half of that of MLCK2(Ig1-4), indicating a significantly greater affinity of MLCK1(Ig1-4) for F-actin (Fig. 6B). However, MLCK1(Ig3) alone bound F-actin poorly (Fig. 6B). Thus, although Ig3 contributes significantly to MLCK1 binding to F-actin, it does so only in concert with surrounding Ig domains.

**Figure 6.**
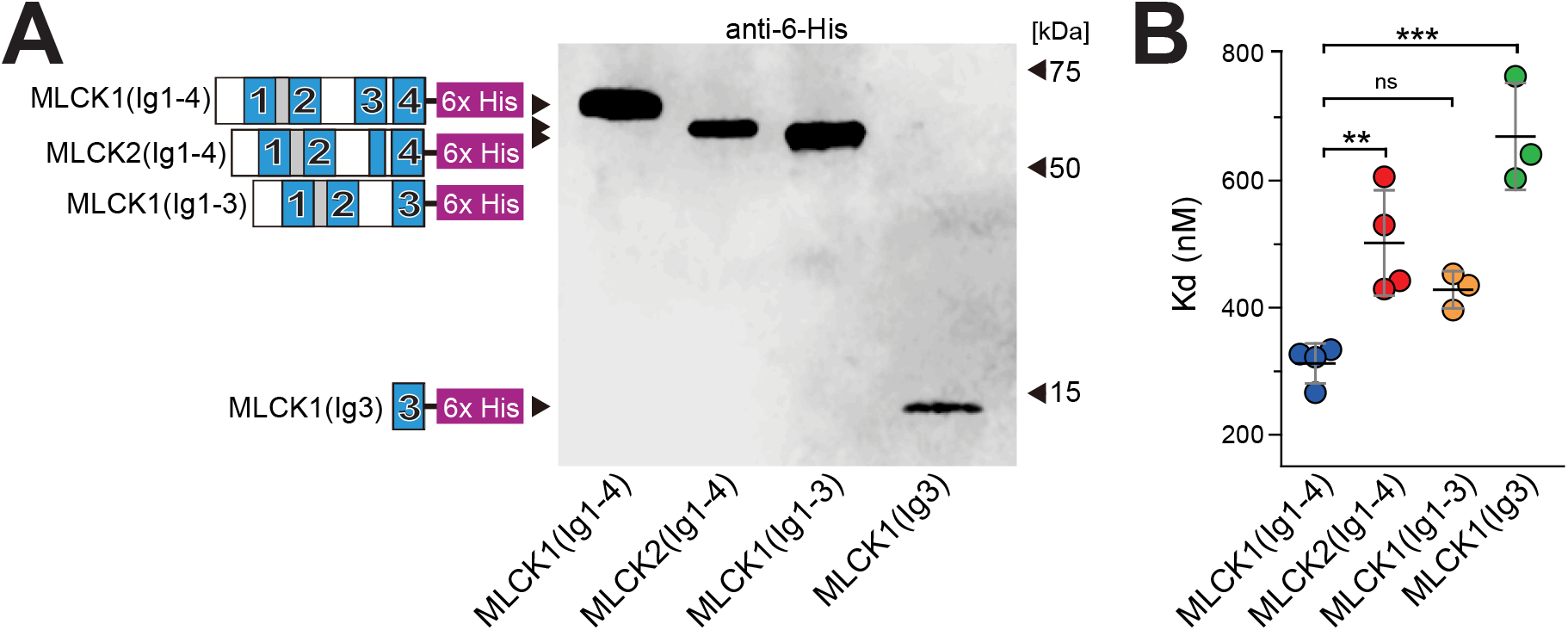
MLCK1 binding affinity for F-actin is greater than MLCK2 binding affinity. **A.** Structures and Western blot of recombinant proteins encoding His-tagged MLCK constructs used for microscale thermophoresis (MST). Ig domains are numbered and shown in blue. **B.** Dissociation constants (Kd) of MLCK protein construct-binding to F-actin. *n* = 3-4. Mean ± SD. ANOVA. **, P <0.01. ***, P <0.001.

### MLCK perijunctional recruitment correlates with F-actin binding affinity

The greater affinity of MLCK1(Ig1-4) for F-actin relative to that of MLCK2(Ig1-4) suggests that this could explain differential recruitment of MLCK1 and MLCK2 to the perijunctional actomyosin ring in polarized epithelial cells. To test this possibility, constructs identical to those studied in vitro as well as MLCK1(Ig2-3) were stably expressed as EGFP fusion proteins in Caco-2_BBe_ cells (Fig. 7A, B). Both MLCK1(Ig1-4) and MLCK1(Ig1-3) were effectively recruited to the perijunctional actomyosin ring (Fig. 7C, D). Perijunctional localization of MLCK2(Ig1-4) and MLCK1(Ig2-3) was detectable, but dramatically reduced relative to MLCK1(Ig1-4) (Fig. 7C, D). Finally, the distribution of MLCK1(Ig3) was similar to that of EGFP alone, which was expressed as a negative control. Thus, the extent to which long MLCK constructs colocalized with perijunctional actomyosin paralleled the affinity of these constructs for F-actin. Differential binding to F-actin may, therefore, explain the distinct steady-state localizations of MLCK1 and MLCK2.

**Figure 7.**
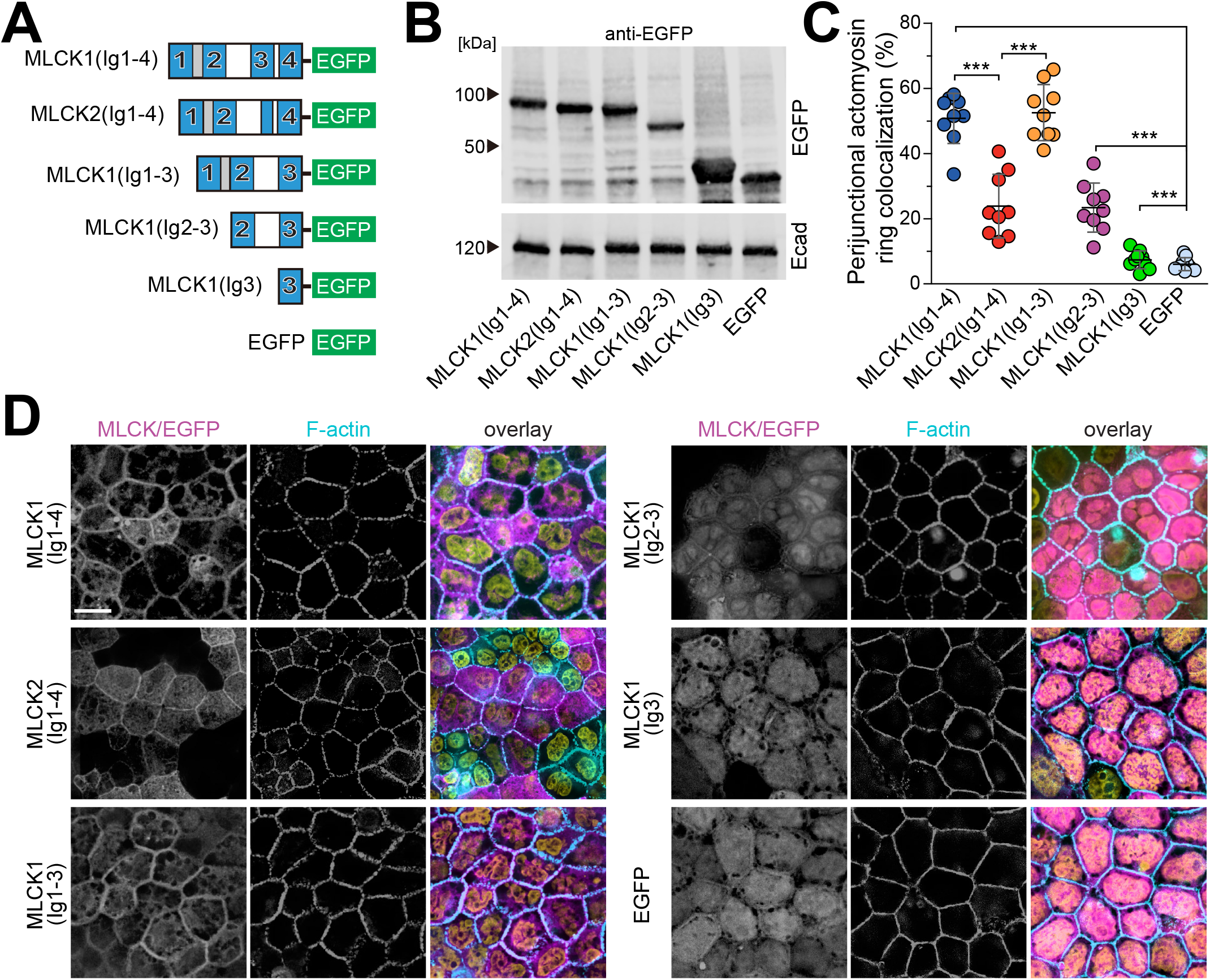
Ig1-3 are required for steady-state MLCK1 recruitment trafficking to the perijunctional actomyosin ring. **A.** Structures of EGFP-tagged MLCK constructs stably expressed in Caco-2_BBe_ cells. Ig domains are numbered and shown in blue. Free EGFP was stably expressed as a control. **B.** Western blot of EGFP-tagged MLCK. **C.** Quantitative morphometry of perijunctional actomyosin ring localization of indicated MLCK constructs or EGFP. *n* = 9. Mean ± SD. ANOVA. ***, P <0.001. **D.** Representative images of EGFP-tagged MLCK constructs or EGFP (magenta), F-actin (cyan), and nuclei (yellow). Bar = 10 μm.

### Ig3 acts as a dominant negative that limits TNF-induced MLCK1 recruitment and barrier loss

Although Ig3 alone was insufficient to direct targeting to the perijunctional actomyosin ring, it was also essential for F-actin binding in vitro and perijunctional recruitment within Caco-2_BBe_ cells. We therefore hypothesized that Ig3 might block critical interactions necessary for steady-state and TNF-induced MLCK1 recruitment. To test this we created Caco-2_BBe_ cells with constitutive and inducible expression of MLCK1-EGFP and mCherry-Ig3, respectively, and mixed these with cells that only expressed MLCK1-EGFP. Cytoplasmic MLCK1 was dramatically increased in cells that also expressed mCherry-Ig3 (Fig. 8A, B). Moreover, TNF-induced MLCK1 recruitment was blocked in cells that coexpressed mCherry-Ig3 (Fig. 8A, B). Ig3-mediated interactions are, therefore, essential for both steady-state and TNF-induced MLCK1 recruitment to the perijunctional actomyosin ring.

**Figure 8.**
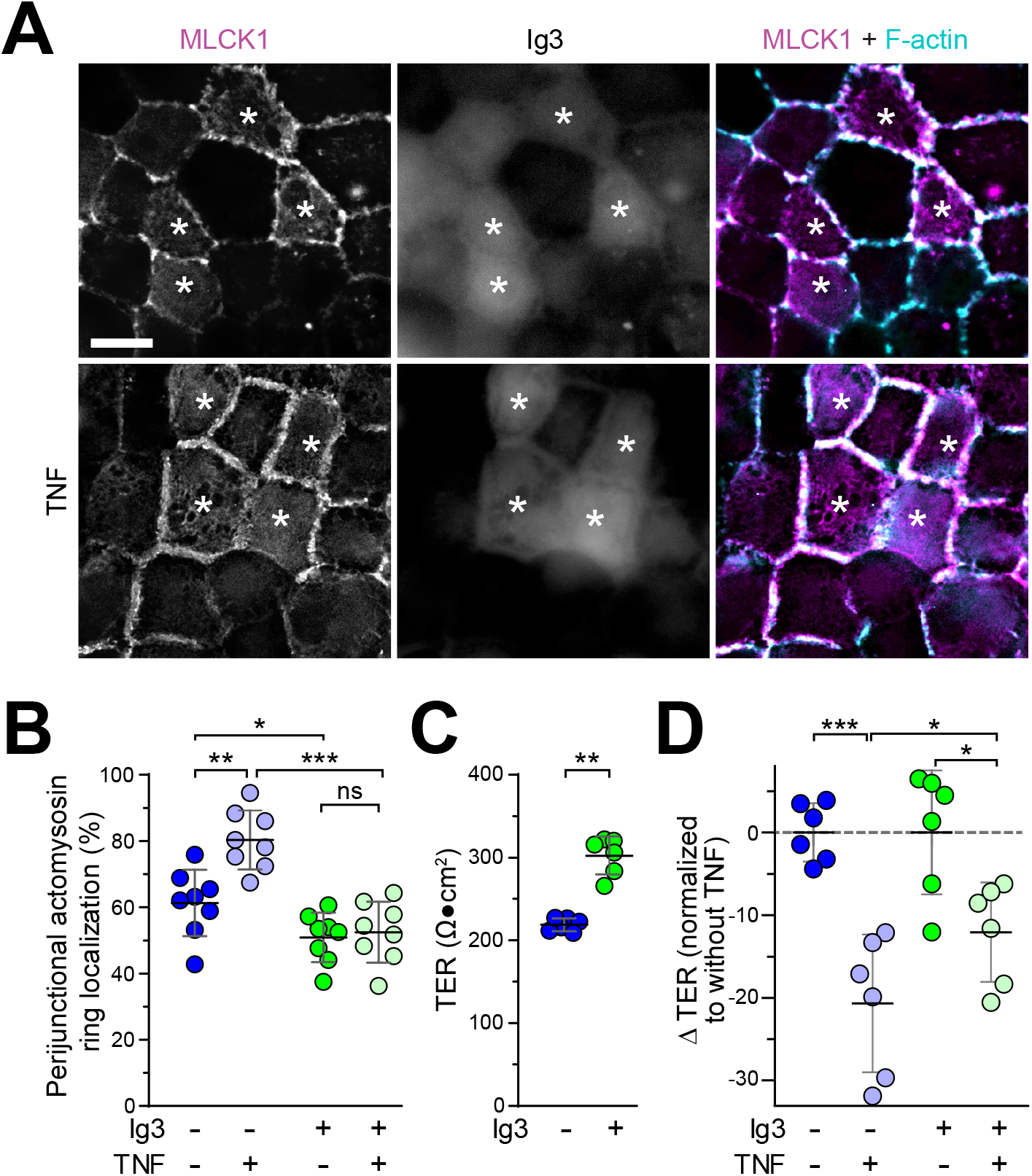
Ig3 acts as a dominant negative to block TNF-induced MLCK1 recruitment to the perijunctional actomyosin ring. **A.** Representative images of Caco-2_BBe_ monolayers expressing MLCK1-EGFP (MLCK1) with and without mCherry-Ig3 expression. Increased cytoplasmic MLCK1 is present in mCherry-Ig3 expressing cells (asterisks), both before and after TNF treatment. The overlay shows MLCK1 (magenta) and F-actin (cyan), but not Ig3. Bar = 10 μm. **B.** Morphometric analysis of perijunctional MLCK1 in the absence (blue symbols) or presence (green symbols) of inducible mCherry-Ig3 expression before (dark symbols) or after (light symbols) TNF treatment. *n* = 8. Mean ± SD. ANOVA. *, P < 0.05. **, P <0.01. ***, P <0.001. **C.** Steady-state transepithelial electrical resistance (TER) of MLCK1-EGFP-expressing monolayers without (blue symbols) or with (green symbols) induction of mCherry-Ig3 expression. *n* = 6. Mean ± SD. Student’s *t* test. **, P < 0.01. **D.** TER of MLCK1-EGFP-expressing monolayers without (blue symbols) or with (green symbols) inducible mCherry-Ig3 expression. TER before TNF treatment is used to normalize TER of similar, i.e., without or with mCherry-Ig3 expression, TNF-treated monolayers in order to facilitate comparison of TNF effects. *n* = 6. Mean ± SD. ANOVA. *, P <0.01, ***, P <0.001.

We next asked whether mCherry-Ig3 expression affected barrier function. At steady-state, mCherry-Ig3 expression-induced a 38%±3% TER increase (Fig. 8C), consistent with MLCK1-EGFP displacement from the perijunctional actomyosin ring (6,18). Although mCherry-Ig3 expression did not completely prevent TNF-induced barrier loss, it did significantly limit the magnitude by which TER was reduced (Fig. 8D). Thus, although Ig3 cannot mediate MLCK1 recruitment and tight junction barrier regulation, it is sufficient to act as a dominant negative and interfere with these processes.

## DISCUSSION

Our previous work using small molecules has demonstrated that TNF-induced MLCK1 recruitment to the perijunctional actomyosin ring has potential as a therapeutic target to prevent inflammation-induced tight junction barrier loss as well as progression of immune-mediated disease (6,18). While these small molecules were selected using an in silico screen for MLCK1 Ig3 binding, the striking results did not exclude the possibility that these small molecules also bound to other regions, including the Ig domains, within MLCK1 (6). Here, we have used a protein-based approach to determine the extent to which Ig3 contributes to MLCK1 recruitment and barrier regulation regions. We also sought to elucidate the mechanisms that direct MLCK1 recruitment. Our data demonstrate that Ig3 alone is insufficient to direct MLCK1 recruitment but is capable of preventing MLCK1 recruitment. Moreover, the correlation with F-actin binding affinity suggests that this may be a significant determinant of perijunctional recruitment.

In addition to analyzing cell culture models and recombinant proteins, we assessed expression and subcellular distributions of MLCK1 and MLCK2 in ileal biopsies from healthy control subjects and Crohn’s disease patients. Quantitative morphometry demonstrated that MLCK1 expression and perijunctional localization were specifically increased in active Crohn’s disease. Although descriptive, these data support the hypothesis that blocking MLCK1 recruitment will be of benefit in Crohn’s disease as well as ulcerative colitis.

We have cloned and characterized the human long MLCK promoter (19). While they explain cytokine-induced upregulation, the processes that regulate long MLCK mRNA splicing to control relative expression of MLCK1 and MLCK2 are undefined. However, human SNPs rs77323602 and rs147245669 have been suggested to favor MLCK1 or MLCK2 expression, respectively, in endothelial cells (32–34). Limited data suggest that this may reflect the ability of heterogeneous nuclear ribonucleoprotein A1 (hnRNPA1) to promote exon 11 removal and, as a direct consequence, production of MLCK2 transcripts (32,33). In the intestine, hnRNPA1 is primarily expressed within undifferentiated epithelia (35), consistent with splicing-induced expression of MLCK2, but not MLCK1, in these cells (17). The hnRNPA1 downregulation that occurs during enterocyte differentiation may, therefore, explain increased MLCK1 transcript abundance and protein expression within villous epithelial. Although not been studied in detail, it is also striking that the Crohn’s disease-associated 3020insC *NOD2* polymorphism has been linked to reduced hnRNPA1 splicing activity (36,37). Thus, although further studies are needed, downregulation of hnRNPA1 activity could be responsible for preferential MLCK1 upregulation in Crohn’s disease. Although it is not possible to develop MLCK2-specific antibodies, we overcame this obstacle by expressing EGFP-tagged MLCK1 and MLCK2 in well-differentiated Caco-2_BBe_ intestinal epithelial cells. This allowed us to recognize the dramatically different subcellular distributions of these splice variants. As previously documented in tissue, MLCK1 was primarily localized to the perijunctional actomyosin ring. This tagged protein approach also allowed us to recognize that MLCK2, which was previously described as diffuse and cytoplasmic, was specifically associated with basal stress fibers and, to a lesser extent, epithelial lateral membranes. Consistent with the recognized importance of cyclic MLC phosphorylation within stress fibers during epithelial migration, MLCK2, but not MLCK1, overexpression inhibited epithelial migration. Long MLCK expressed within endothelial cells is primarily MLCK2 (34). Together with the demonstrated contributions of stress fiber contraction to endothelial barrier regulation (24,38-40), these data support the idea that MLCK2 primarily regulates microfilaments while, at least in intestinal epithelia, MLCK1 is the principal regulator of perijunctional actomyosin and tight junction permeability (23-25,41). Overall, these data suggest that the two long MLCK splice variants expressed in intestinal epithelium preferentially associate with and modulate different actin-rich structures to have distinct effects on epithelial function.

The striking differences between MLCK1 and MLCK2 are surprising given that these proteins differ by only 69 amino acids that comprise less than 4% of the primary sequence. Nevertheless, this sequence completes the Ig3 domain within the N-terminal extension that differentiates long from short MLCK proteins that are products of a single *MYLK* gene (34,42). We therefore focused on the N-terminal extension to define the unique behaviors of MLCK1 and MLCK2. First, we showed that the Ig1-4 region of MLCK1, but not MLCK2, is sufficient to direct steady-state and TNF-induced recruitment to the perijunctional actomyosin ring. In addition to restricting the region that directs MLCK1 recruitment, these data indicate that enzymatic activity, is not required for MLCK1 recruitment.

Previous studies have identified diverse functions for the Ig domains present within long MLCK, including recruitment to actin rich structures and in vitro binding to microfilaments (28,31,43,44). Ig1-2 was reported to bind to F-actin in vitro with a Kd of 0.39 μM (31), similar to the ∼0.50 μM Kd we measured for MLCK2(Ig1-4) but somewhat greater than the 0.31 μM Kd we measured for MLCK1(Ig1-4). Finally, our data indicate that the five F-actin-binding DXR motifs (28,45) present within both MLCK1 and MLCK2 are not required for recruitment to the perijunctional actomyosin ring.

Despite binding F-actin poorly and failing to direct recruitment to the perijunctional actomyosin ring, Ig3 was an effective dominant negative inhibitor that displaced MLCK1 from the perijunctional actomyosin ring, increased steady-state barrier function, blocked TNF-induced MLCK1 recruitment, and limited TNF-induced barrier loss. This is consistent with our recent discovery that FKBP8 is a chaperone protein that binds MLCK1(Ig1-4) and is essential for TNF - induced MLCK1 recruitment and TNF-induced barrier loss (6,18). Thus, while not sufficient, Ig3 is necessary for TNF-induced, MLCK1-mediated tight junction regulation.

As a whole, our data indicate that MLCK1 is preferentially upregulated in human Crohn’s disease, where it is most concentrated within the perijunctional actomyosin ring of intestinal epithelial cells. Together with our definition of the domains required for MLCK1 recruitment to the perijunctional actomyosin ring and demonstration that Ig3 plays a critical role in this process, these data create a foundation that will allow development of MLCK1(Ig1-4)- and Ig3-targeted molecular therapeutic approaches.

## MATERIALS AND METHODS

### Human tissues

De-identified ileal biopsies from Crohn’s disease patients were obtained from Brigham and Women’s Hospital under IRB-approved protocols. Only well-oriented ileal biopsies with excellent tissue preservation were used. Biopsies with extensive ulceration were excluded. De-identified Ileal intestinal biopsies without abnormalities from age- and sex-matched healthy patients, e.g., undergoing screening colonoscopies, were used as controls. Formalin-fixed, paraffin-embedded biopsies were assembled into tissue microarrays prior to staining.

### Cell lines

Plasmids with expression cassettes flanked by transposable elements were co-transfected with piggyBac transposase (System Biosciences) into clone 5E6L Caco-2_BBe_ cells. Expression cassettes encoded EGFP- or FR-tagged full length MLCK1 or MLCK2 as well as indicated mutants. Separate populations were stably-transfected to express MLCK1-EGFP constitutively and mCherry-Ig3 inducibly via a Tet-on3G system (Clontech). Cells were maintained in high-glucose Dulbecco’s modified Eagle’s medium (Life Technologies) supplemented with 10% fetal calf serum, and 15 mM HEPES, pH 7.4, as previously-described (18,46). Cells were treated with doxycycline (20 ng/ml) to induce mCherry-Ig3 expression.

For growth as polarized monolayers, Transwell inserts (Corning) were coated with rat tail collagen and plated as described (47). Cytokines were added basolaterally, with IFN-γ added 18 hours before TNF, which was added for 4 hours, as described (47).

For migration assays, cells were plated in 2 well inserts in µ-dish 35 mm plates (Ibidi). Inserts were removed to allow for migration over a defined area after cells within the wells became confluent.

### Antibodies

Antigen affinity-purified rabbit anti-E-cadherin (Cell Signaling Technology, AB_2291471) was used for western blot (0.1 µg/ml); antigen affinity-purified rabbit anti-GFP (Novus, AB_523903) was used for western blot (2.2 µg/ml); antigen affinity-purified rabbit anti-long MLCK (AB_2921292) was used for immunofluorescence (10 µg/ml) and western blot (2.5 µg/ml); mouse anti-6-His (Bio X Cell, AB_2687790) was used for western blot (3 µg/ml); antigen affinity-purified rat anti-MLCK1 (AB_2909525) was used for immunofluorescence (20 µg/ml); monoclonal rat anti-ZO-1 (AB_2783858) was used for immunofluorescence (1 µg/ml); and mouse monoclonal anti-γ-actin (Santa Cruz, AB_2890619) was used for immunofluorescence (0.4 µg/ml). Secondary antibodies conjugated to IRDye 680LT or IRDye 800CW (Li-Cor) were used for western blot and highly cross-adsorbed IgG F(ab′)_2_ secondary antibodies conjugated to Alexafluor 594 or Alexafluor 647 (Jackson Immunoresearch) were used for immunofluorescence.

### Latrunculin A treatment and cytosolic extraction

Caco-2_BBe_ monolayers stably expressing EGFP-tagged MLCK1 and MLCK2 were equilibrated in HBSS for 30 min at 37 °C before the treatment. Then, the basolateral media was replaced by HBSS containing 1 μM LatA to allow actin depolymerization and cells were incubated at 37 °C (29). Monolayers were fixed before or 30 min after LatA treatment, stained for F-actin using Alexafluor 594 phalloidin, and imaged.

For extraction, cells were washed once with HBSS which was then removed and gently replaced with extraction buffer (50 mM MES, 3 mM EGTA, 5 mM MgCl_2_, 0.5% Triton X-100, p H7.4), as described (48). Images were collected at time 0, and 2 and 4 min after CSK buffer treatment.

### Electrophysiology

Monolayers were transferred from culture medium to HBSS supplemented with 15 mM HEPES, pH 7.4, and 25 mM D-glucose. After equilibration, potential differences were measured before and during application of a 50-μA current. Transepithelial short-circuit current (Isc) and transepithelial resistance (TER) were calculated using Ohm’s law, as previously described (2,49).

### Immunostains

Formalin-fixed, paraffin-embedded ileal samples from age-sex matched Crohn’s patients and healthy controls were assembled into tissue microarrays. Sections were deparaffinized and rehydrated and underwent antigen retrieval using Trilogy (Sigma-Aldrich) solution and a pressure cooker. After cooling, sections were incubated in bleaching buffer under a bright LED (broad spectrum) light for 1 hour at room temperature (50). After replenishing with fresh bleaching buffer, incubation under light was continued for an additional 1 hour. After incubation with blocking buffer (Agilent), primary antibodies prepared in antibody diluent (Agilent) were added for 8 hours at room temperature. After 5 washes, secondary antibodies with Hoechst 33342 (1 μg/ml) were added for 1 hour. After washing, sections were mounted using Vectashield Plus Antifade Medium (Vector Laboratories).

Caco-2_BBe_ cells were fixed in 1% paraformaldehyde-lysine-periodate fixative (51) and immunostained as described previously using the antibodies described above. F-actin was labeled using Alexafluor 594- or Alexafluor 680-conjugated phalloidin.

### Imaging

Widefield imaging was performed using an Axioplan 2 widefield microscope (Zeiss) with 20x NA 0.8 Plan-Apochromat and 100x NA 1.4 Plan-Neofluar objectives, ET filter cubes (Chroma), LED illuminator (Lumencor SOLA), and Coolsnap HQ (Photometrics).

Three systems were used for acquisition of confocal images. A CSU-X1 spinning disk (Yokogawa) system mounted on a DMI6000 microscope (Leica) with 100x NA 1.4 HC PLAN APO CS2 objective, 4-channel solid-state Spectral-Borealis laser launch (Andor), a Zyla 4.2 Plus sCMOS camera (Andor) A STEDYCON (Abberior) system mounted on a Zeiss Axio observer Z1 microscope equipped with a 100X NA 1.46 α Plan-Apochromat objective (Zeiss) was used in laserscanning confocal mode. Finally, a Zeiss LSM 880 with Airyscan equipped with a 63X/ NA 1.40 Plan-Apochromat objective (Zeiss) was used in super resolution mode with 0.16 μm/pixel, 1024 x 1024 frame size, 2x averaging, and 4x digital zoom.

Tiled tissue microarray images (widefield) were collected using a DM4000 widefield microscope with 20X NA 0.7 HC PLAN APO objective (Leica), ORCA-Flash 4.0 LT+ CMOS camera (Hamamatsu), motorized emission filter wheel and xyz stage (Ludl), 5 channel Aura light engine (Lumencor), and multichannel dichroic matched to single band excitation filters (Semrock), as described (50).

### Post-acquisition analysis and morphometry

Post-acquisition processing and analysis used MetaMorph 7 (Molecular Devices), Autoquant X3 (MediaCybernetics), FIJI (52), and Imaris 9.9.0 (Bitplane). For MLCK1 and total MLCK analysis in human biopsies, quantitative morphometry was performed using Cell Profiler (53). Villus and crypt regional masks with whole biopsy tiled scans were created manually, and whole epithelial and apical epithelial masks were generated based on γ-actin distribution. Intensities were normalized to γ-actin after preliminary validation showing that γ-actin expression was unchanged in Crohn’s disease.

For analysis of cultured cells, the perijunctional actomyosin ring was defined based on phalloidin staining. ImageJ (National Institutes of Health, Bethesda, MD, USA) was used to create a region of interest (ROI) around the complete cell and a discrete intracellular ROI. The larger ROI was taken as total fluorescence. The fluorescence of the smaller ROI was subtracted from total and then normalized to total fluorescence to determine the perijunctional MLCK fraction (6). For analysis of stress fiber area, Zeiss Airyscan images were rendered using IMARIS 9.9.0. The Imaris surface function was used in tandem with the machine learning classification and the FIJI Labkit plugin.

### Recombinant protein preparation, actin polymerization, and microscale thermophoresis

His-tagged recombinant proteins were expressed in Rosetta2 (DE3)pLysS bacteria (Novagen) and purified on IMAC columns (Bio-Rad) using an AktaPure FPLC system (Cytiva). Purity was assessed by SDS-PAGE.

Actin binding and polymerization were assessed using the actin binding protein kit (Cytoskeleton, BK013). Actin pellet gel bands were quantified by densitometry using FIJI.

In vitro actin polymerization was assessed using the actin polymerization kit (Cytoskeleton, BK003). Briefly, 60 µL of 4.53 μM pyrene-labelled G-actin was polymerized in presence or absence of 2 μM MLCK1(Ig1-4) or MLCK2(Ig1-4). Fluorescence was monitored using a Synergy H1 (Biotek) plate reader at excitation and emission wavelengths of 365 and 407 nm, respectively.

His-tagged MLCK proteins were labeled using a His-Tag Labeling Kit RED-tris-NTA (NanoTemper). Microscale thermophoresis was performed using a Monolith NT.115 (NanoTemper). Data were analyzed by MO Affinity Analysis software. F-actin was polymerized from chicken gizzard (smooth muscle) muscle actin (Cytoskeleton, APHL99).

### Statistics

All data are presented as mean ± SD and are representative of at least 3 independent experiments. Statistical significance was determined by 2-tailed Student’s *t* test (unpaired) or 1-way ANOVA with Bonferroni’s correction, as indicated in each figure legend using GraphPad Prism 9. *, P < 0.05; **, P < 0.01; ***, P < 0.001.

## Acknowledgments

We thank Drs. W Vallen Graham, Yingmin Wang, Yi-Tang Wang, and Harry Rosenberg for their contributions to early phases of this work, Dr. Nitesh Shashikanth and Dr. Jose Yeste for their assistance with confocal microscopy and image processing, Dr. Kelly L. Arnett and the Harvard Center for Macromolecular Interactions for instruction and assistance with microscale thermophoresis, Dr. Lay-Hong Ang for instruction and assistance with Airyscan microscope, and Heather Marlett (Nationwide Histology) for outstanding tissue preparation.

## Author contributions

Conceptualization: SC-P, WH, JRT. Experimentation: SC-P, WH. Data analysis SC-P, WH, SA, JZ, LZ, JRT. Manuscript preparation: SC-P, WH, JRT. Manuscript editing: SC-P, WH, SA, JZ, LZ, JRT.

## Data sharing

Data that support the findings of this study are included in the main text. Requests for additional information or materials may be addressed to the corresponding author.

## REFERENCES

1. Zolotarevsky, Y., Hecht, G., Koutsouris, A., Gonzalez, D. E., Quan, C., Tom, J., Mrsny, R. J., and Turner, J. R. (2002) A membrane-permeant peptide that inhibits MLC kinase restores barrier function in in vitro models of intestinal disease. Gastroenterol. 123, 163–172

2. Turner, J. R., Rill, B. K., Carlson, S. L., Carnes, D., Kerner, R., Mrsny, R. J., and Madara, J. L. (1997) Physiological regulation of epithelial tight junctions is associated with myosin light-chain phosphorylation. Am. J. Physiol. 273, C1378–1385

3. Clayburgh, D. R., Barrett, T. A., Tang, Y., Meddings, J. B., Van Eldik, L. J., Watterson, D. M., Clarke, L. L., Mrsny, R. J., and Turner, J. R. (2005) Epithelial myosin light chain kinase-dependent barrier dysfunction mediates T cell activation-induced diarrhea in vivo. J. Clin. Invest. 115, 2702–2715

4. Su, L., Nalle, S. C., Shen, L., Turner, E. S., Singh, G., Breskin, L. A., Khramtsova, E. A., Khramtsova, G., Tsai, P. Y., Fu, Y. X., Abraham, C., and Turner, J. R. (2013) TNFR2 activates MLCK-dependent tight junction dysregulation to cause apoptosis-mediated barrier loss and experimental colitis. Gastroenterol. 145, 407–415

5. Su, L., Shen, L., Clayburgh, D. R., Nalle, S. C., Sullivan, E. A., Meddings, J. B., Abraham, C., and Turner, J. R. (2009) Targeted epithelial tight junction dysfunction causes immune activation and contributes to development of experimental colitis. Gastroenterol. 136, 551–563

6. Graham, W. V., He, W., Marchiando, A. M., Zha, J., Singh, G., Li, H. S., Biswas, A., Ong, M., Jiang, Z. H., Choi, W., Zuccola, H., Wang, Y., Griffith, J., Wu, J., Rosenberg, H. J., Wang, Y., Snapper, S. B., Ostrov, D., Meredith, S. C., Miller, L. W., and Turner, J. R. (2019) Intracellular MLCK1 diversion reverses barrier loss to restore mucosal homeostasis. Nat. Med. 25, 690–700

7. Ma, T. Y., Boivin, M. A., Ye, D., Pedram, A., and Said, H. M. (2005) Mechanism of TNF-{alpha} modulation of Caco-2 intestinal epithelial tight junction barrier: role of myosin light-chain kinase protein expression. Am. J. Physiol. - Gastrointest. Liver Physiol. 288, G422–430

8. Wang, F., Graham, W. V., Wang, Y., Witkowski, E. D., Schwarz, B. T., and Turner, J. R. (2005) Interferon-gamma and tumor necrosis factor-alpha synergize to induce intestinal epithelial barrier dysfunction by up-regulating myosin light chain kinase expression. Am. J. Pathol. 166, 409–419

9. Al-Sadi, R., Ye, D., Dokladny, K., and Ma, T. Y. (2008) Mechanism of IL-1beta-induced increase in intestinal epithelial tight junction permeability. J. Immunol. 180, 5653–5661

10. Feighery, L. M., Cochrane, S. W., Quinn, T., Baird, A. W., O’Toole, D., Owens, S. E., O’Donoghue, D., Mrsny, R. J., and Brayden, D. J. (2008) Myosin light chain kinase inhibition: correction of increased intestinal epithelial permeability in vitro. Pharm. Res. 25, 1377-1386

11. Chen, C., Wang, P., Su, Q., Wang, S., and Wang, F. (2012) Myosin light chain kinase mediates intestinal barrier disruption following burn injury. PLoS One 7, e34946

12. Guo, M., Yuan, S. Y., Frederich, B. J., Sun, C., Shen, Q., McLean, D. L., and Wu, M. H. (2012) Role of non-muscle myosin light chain kinase in neutrophil-mediated intestinal barrier dysfunction during thermal injury. Shock 38, 436–443

13. Zahs, A., Bird, M. D., Ramirez, L., Turner, J. R., Choudhry, M. A., and Kovacs, E. J. (2012) Inhibition of long myosin light-chain kinase activation alleviates intestinal damage after binge ethanol exposure and burn injury. Am. J. Physiol. - Gastrointest. Liver Physiol. 303, G705–712

14. Lorentz, C. A., Liang, Z., Meng, M., Chen, C. W., Yoseph, B. P., Breed, E. R., Mittal, R., Klingensmith, N. J., Farris, A. B., Burd, E. M., Koval, M., Ford, M. L., and Coopersmith, C. M. (2017) Myosin light chain kinase knockout improves gut barrier function and confers a survival advantage in polymicrobial sepsis. Mol Med 23, 155–165

15. Nalle, S. C., Zuo, L., Ong, M., Singh, G., Worthylake, A. M., Choi, W., Manresa, M. C., Southworth, A. P., Edelblum, K. L., Baker, G. J., Joseph, N. E., Savage, P. A., and Turner, J. R. (2019) Graft-versus-host disease propagation depends on increased intestinal epithelial tight junction permeability. J. Clin. Invest. 129, 902–914

16. Pai, Y. C., Weng, L. T., Wei, S. C., Wu, L. L., Shih, D. Q., Targan, S. R., Turner, J. R., and Yu, L. C. (2020) Gut microbial transcytosis induced by tumor necrosis factor-like 1A-dependent activation of a myosin light chain kinase splice variant contributes to IBD. Journal of Crohn’s & colitis 15, 258–272

17. Clayburgh, D. R., Rosen, S., Witkowski, E. D., Wang, F., Blair, S., Dudek, S., Garcia, J. G., Alverdy, J. C., and Turner, J. R. (2004) A differentiation-dependent splice variant of myosin light chain kinase, MLCK1, regulates epithelial tight junction permeability. J. Biol. Chem. 279, 55506-55513

18. Zuo, L., Kuo, W. T., Cao, F., Chanez-Paredes, S. D., Zeve, D., Mannam, P., Jean-Francois, L., Day, A., Vallen Graham, W., Sweat, Y. Y., Shashikanth, N., Breault, D. T., and Turner, J. R. (2023) Tacrolimus-binding protein FKBP8 directs myosin light chain kinase-dependent barrier regulation and is a potential therapeutic target in Crohn’s disease. Gut 72, 870–881

19. Graham, W. V., Wang, F., Clayburgh, D. R., Cheng, J. X., Yoon, B., Wang, Y., Lin, A., and Turner, J. R. (2006) Tumor necrosis factor-induced long myosin light chain kinase transcription is regulated by differentiation-dependent signaling events. Characterization of the human long myosin light chain kinase promoter. J. Biol. Chem. 281, 26205–26215

20. Lukas, T. J., Mirzoeva, S., Slomczynska, U., and Watterson, D. M. (1999) Identification of novel classes of protein kinase inhibitors using combinatorial peptide chemistry based on functional genomics knowledge. J. Med. Chem. 42, 910–919

21. Owens, S. E., Graham, W. V., Siccardi, D., Turner, J. R., and Mrsny, R. J. (2005) A strategy to identify stable membrane-permeant peptide inhibitors of myosin light chain kinase. Pharm. Res. 22, 703–709

22. Chen, X., Pavlish, K., and Benoit, J. N. (2008) Myosin phosphorylation triggers actin polymerization in vascular smooth muscle. Am. J. Physiol. - Heart Circ. Physiol. 295, H2172–2177

23. Shen, L., Black, E. D., Witkowski, E. D., Lencer, W. I., Guerriero, V., Schneeberger, E. E., and Turner, J. R. (2006) Myosin light chain phosphorylation regulates barrier function by remodeling tight junction structure. J. Cell Sci. 119, 2095–2106

24. Goeckeler, Z. M., and Wysolmerski, R. B. (1995) Myosin light chain kinase-regulated endothelial cell contraction: the relationship between isometric tension, actin polymerization, and myosin phosphorylation. J Cell Biol 130, 613–627

25. Katoh, K., Kano, Y., Amano, M., Kaibuchi, K., and Fujiwara, K. (2001) Stress fiber organization regulated by MLCK and Rho-kinase in cultured human fibroblasts. Am. J. Physiol. - Cell Physiol. 280, C1669–1679

26. Schmidt, J. T., Morgan, P., Dowell, N., and Leu, B. (2002) Myosin light chain phosphorylation and growth cone motility. J. Neurobiol. 52, 175–188

27. Chen, C., Tao, T., Wen, C., He, W. Q., Qiao, Y. N., Gao, Y. Q., Chen, X., Wang, P., Chen, C. P., Zhao, W., Chen, H. Q., Ye, A. P., Peng, Y. J., and Zhu, M. S. (2014) Myosin light chain kinase (MLCK) regulates cell migration in a myosin regulatory light chain phosphorylation-independent mechanism. J. Biol. Chem. 289, 28478–28488

28. Smith, L., Parizi-Robinson, M., Zhu, M. S., Zhi, G., Fukui, R., Kamm, K. E., and Stull, J. T. (2002) Properties of long myosin light chain kinase binding to F-actin in vitro and in vivo. J. Biol. Chem. 277, 35597–35604

29. Shen, L., and Turner, J. R. (2005) Actin depolymerization disrupts tight junctions via caveolae-mediated endocytosis. Mol. Biol. Cell 16, 3919–3936

30. Birukov, K. G., Csortos, C., Marzilli, L., Dudek, S., Ma, S. F., Bresnick, A. R., Verin, A. D., Cotter, R. J., and Garcia, J. G. (2001) Differential regulation of alternatively spliced endothelial cell myosin light chain kinase isoforms by p60(Src). J. Biol. Chem. 276, 8567–8573

31. Yang, C. X., Chen, H. Q., Chen, C., Yu, W. P., Zhang, W. C., Peng, Y. J., He, W. Q., Wei, D. M., Gao, X., and Zhu, M. S. (2006) Microfilament-binding properties of N-terminal extension of the isoform of smooth muscle long myosin light chain kinase. Cell Res 16, 367–376

32. Mascarenhas, J. B., Tchourbanov, A. Y., Fan, H., Danilov, S. M., Wang, T., and Garcia, J. G. (2017) Mechanical Stress and Single Nucleotide Variants Regulate Alternative Splicing of the MYLK Gene. Am. J. Respir. Cell Mol. Biol. 56, 29–37

33. Mascarenhas, J. B., Tchourbanov, A. Y., Danilov, S. M., Zhou, T., Wang, T., and Garcia, J. G. N. (2018) The Splicing Factor hnRNPA1 Regulates Alternate Splicing of the MYLK Gene. Am. J. Respir. Cell Mol. Biol. 58, 604–613

34. Lazar, V., and Garcia, J. G. (1999) A single human myosin light chain kinase gene (MLCK; MYLK). Genomics 57, 256–267

35. Karlsson, M., Zhang, C., Mear, L., Zhong, W., Digre, A., Katona, B., Sjostedt, E., Butler, L., Odeberg, J., Dusart, P., Edfors, F., Oksvold, P., von Feilitzen, K., Zwahlen, M., Arif, M., Altay, O., Li, X., Ozcan, M., Mardinoglu, A., Fagerberg, L., Mulder, J., Luo, Y., Ponten, F., Uhlen, M., and Lindskog, C. (2021) A single-cell type transcriptomics map of human tissues. Sci Adv 7

36. Noguchi, E., Homma, Y., Kang, X., Netea, M. G., and Ma, X. (2009) A Crohn’s disease-associated NOD2 mutation suppresses transcription of human IL10 by inhibiting activity of the nuclear ribonucleoprotein hnRNP-A1. Nat. Immunol. 10, 471–479

37. Clarke, J. P., Thibault, P. A., Salapa, H. E., and Levin, M. C. (2021) A Comprehensive Analysis of the Role of hnRNP A1 Function and Dysfunction in the Pathogenesis of Neurodegenerative Disease. Front Mol Biosci 8, 659610

38. Garcia, J. G., Davis, H. W., and Patterson, C. E. (1995) Regulation of endothelial cell gap formation and barrier dysfunction: role of myosin light chain phosphorylation. J. Cell. Physiol. 163, 510–522

39. Vandenbroucke, E., Mehta, D., Minshall, R., and Malik, A. B. (2008) Regulation of endothelial junctional permeability. Ann. N. Y. Acad. Sci. 1123, 134–145

40. Jambusaria, A., Hong, Z., Zhang, L., Srivastava, S., Jana, A., Toth, P. T., Dai, Y., Malik, A. B., and Rehman, J. (2020) Endothelial heterogeneity across distinct vascular beds during homeostasis and inflammation. eLife 9

41. Turner, J. R., and Basson, M. D. (2002) Roles of the extracellular matrix and cytoskeleton in intestinal epithelial restitution. in Gastrointestinal Mucosal Repair and Experimental Therapeutics (Hob, C. H., and Wang, J. Y. eds.), Karger Press. pp 1-13

42. Kamm, K. E., and Stull, J. T. (2001) Dedicated myosin light chain kinases with diverse cellular functions. J. Biol. Chem. 276, 4527–4530

43. Dulyaninova, N. G., Patskovsky, Y. V., and Bresnick, A. R. (2004) The N-terminus of the long MLCK induces a disruption in normal spindle morphology and metaphase arrest. J. Cell Sci. 117, 1481–1493

44. Zhang, W. C., Peng, Y. J., He, W. Q., Lv, N., Chen, C., Zhi, G., Chen, H. Q., and Zhu, M. S. (2008) Identification and functional characterization of an aggregation domain in long myosin light chain kinase. FEBS J 275, 2489–2500

45. Poperechnaya, A., Varlamova, O., Lin, P. J., Stull, J. T., and Bresnick, A. R. (2000) Localization and activity of myosin light chain kinase isoforms during the cell cycle. J Cell Biol 151, 697–708

46. Buschmann, M. M., Shen, L., Rajapakse, H., Raleigh, D. R., Wang, Y., Wang, Y., Lingaraju, A., Zha, J., Abbott, E., McAuley, E. M., Breskin, L. A., Wu, L., Anderson, K., Turner, J. R., and Weber, C. R. (2013) Occludin OCEL-domain interactions are required for maintenance and regulation of the tight junction barrier to macromolecular flux. Mol. Biol. Cell 24, 3056–3068

47. Pongkorpsakol, P., Turner, J. R., and Zuo, L. (2020) Culture of Intestinal Epithelial Cell Monolayers and Their Use in Multiplex Macromolecular Permeability Assays for In Vitro Analysis of Tight Junction Size Selectivity. Curr Protoc Immunol 131, e112

48. Zhao, H., Shiue, H., Palkon, S., Wang, Y., Cullinan, P., Burkhardt, J. K., Musch, M. W., Chang, E. B., and Turner, J. R. (2004) Ezrin regulates NHE3 translocation and activation after Na+-glucose cotransport. Proc. Natl. Acad. Sci. U.S.A. 101, 9485–9490

49. Shashikanth, N., Rizzo, H. E., Pongkorpsakol, P., Heneghan, J. F., and Turner, J. R. (2021) Electrophysiologic Analysis of Tight Junction Size and Charge Selectivity. Curr Protoc 1, e143

50. Abtahi, S., Gliksman, N. R., Heneghan, J. F., Nilsen, S. P., Muhlich, J. L., Copeland, J., Rozbicki, E., Allan, C., Dudeja, P. K., and Turner, J. R. (2021) A simple method for creating a high-content microscope for imaging multiplexed tissue microarrays. Curr Protoc 1, e68

51. McLean, I. W., and Nakane, P. K. (1974) Periodate-lysine-paraformaldehyde fixative. A new fixation for immunoelectron microscopy. J. Histochem. Cytochem. 22, 1077–1083

52. Schindelin, J., Arganda-Carreras, I., Frise, E., Kaynig, V., Longair, M., Pietzsch, T., Preibisch, S., Rueden, C., Saalfeld, S., Schmid, B., Tinevez, J. Y., White, D. J., Hartenstein, V., Eliceiri, K., Tomancak, P., and Cardona, A. (2012) Fiji: an open-source platform for biological-image analysis. Nat Methods 9, 676-682

53. McQuin, C., Goodman, A., Chernyshev, V., Kamentsky, L., Cimini, B. A., Karhohs, K. W., Doan, M., Ding, L., Rafelski, S. M., Thirstrup, D., Wiegraebe, W., Singh, S., Becker, T., Caicedo, J. C., and Carpenter, A. E. (2018) CellProfiler 3.0: Next-generation image processing for biology. PLoS Biol 16, e2005970

